# Linking metastatic potential and viscoelastic properties of breast cancer spheroids via dynamic compression and relaxation in microfluidics

**DOI:** 10.1101/2024.07.23.604808

**Authors:** Margherita Tavasso, Ankur D. Bordoloi, Elsa Tanré, Sanne A. H. Dekker, Valeria Garbin, Pouyan E. Boukany

**Affiliations:** Department of Chemical Engineering, Delft University of Technology, Delft, The Netherlands; École Polytechnique, Institut Polytechnique de Paris, Palaiseau, France

**Keywords:** Tissue mechanics, microfluidics, spheroid viscoelasticity, deformation and relaxation, malignancy

## Abstract

The growth and invasion of solid tumors are associated with changes in their viscoelastic properties, influenced by both internal cellular factors and physical forces in the tumor microenvironment. Due to the lack of a comprehensive investigation of tumor tissue viscoelasticity, the relationship between such physical properties and cancer malignancy remains poorly understood. Here, the viscoelastic properties of breast cancer spheroids, 3D (*in vitro*) tumor models, are studied in relation to their metastatic potentials by imposing controlled, dynamic compression within a microfluidic constriction, and subsequently monitoring the relaxation of the imposed deformation. By adopting a modified Maxwell model to extract viscoelastic properties from the compression data, the benign (MCF-10A) spheroids are found to have higher bulk elastic modulus and viscosity compared to malignant spheroids (MCF-7 and MDA-MB-231). The relaxation is characterized by two timescales, captured by a double exponential fitting function, which reveals a similar fast rebound for MCF-7 and MCF-10A. Both the malignant spheroids exhibit similar long-term relaxation and display residual deformation. However, they differ significantly in morphology, particularly in intercellular movements. These differences between malignant spheroids are demonstrated to be linked to their cytoskeletal organization, by microscopic imaging of F-actin within the spheroids, together with cell-cell adhesion strength.

## 1 Introduction

The progression and invasion of solid tumors involve a complex interplay between tumor cells and their surrounding environment including blood and lymph vessels, immune cells, fibroblasts and extracellular matrix (ECM) [1, 2]. From a physical perspective, such interactions translate into solid stress (compressive and tensile forces applied by non-fluidic elements of the tumor environment) and interstitial fluid pressure, arising from tumor growth in confined spaces, ECM stiffening and remodeling, leaky blood vessels and altered lymphatic drainage system [3]. The mechanical properties of tissues, such as elasticity and viscosity, are significantly influenced by these cues, and instigate biochemical changes, which ultimately facilitate metastasis [4, 5].

The viscoelastic characterization of tumor cells has become a key indicator of tumor development and metastasis [6]. Nanomechanical analysis, performed using atomic force microscopy (AFM) of individual metastatic cells (from breast, lung and pancreatic cancer) in pleural fluids of patients showed that cancer cells are softer than their healthy counterparts [7]. This cell softening serves as a biomechanical adaptation to facilitate cancer invasion [8]. Several studies highlighted how confinement and compression can alter the intracellular structure of single cancer cells, linking their mechanical properties to their metastatic potential at the single-cell level. Malignant breast cancer cells traveling through microchannels that mimic blood vessels demonstrated to exhibit greater deformability compared to benign cell types [9, 10]. Additionally, invasive breast cancer cells showed residual irreversible deformations after squeezing through a tight constricted microchannel under flow conditions, attributed to cytoskeleton rearrangements [11]. Furthermore, nuclear envelope rupture and DNA damage occurred in several cancer cell lines during confined migration through narrow spaces and external compression [12].

At the tissue level, tumor cells are densely packed, held together by cell-cell junctions and surrounded by denser extracellular matrix, resulting in a stiffer tumor tissue compared to the surrounding healthy tissue [13, 14]. For this reason, numerous studies investigated how the mechanical properties of the ECM can profoundly alter single and collective cell migration from a primary tumor: the response to different levels of compressive, tensile and elongational stresses can be monitored in *in vitro* 3D tumor models (spheroids) by tuning the viscoelasticity, stiffness and composition of the surrounding matrix [15, 16, 17, 18]. Theseapproaches, however, only analyzed the solid tumor response in static conditions and in relation to the ECM composition [16], making it difficult to identify the intrinsic spheroid viscoelastic properties in response to dynamic external stresses. Solid and interstitial stress vary over a range of 0.21-20 kPa [19, 20] depending on the tissue examined and the tumor stage. Real time monitoring of 3D *in vitro* assays that impose and release controlled stress under biologically relevant conditions can unravel these properties.

Microfluidics provides a powerful platform to explore the viscoelastic properties of cancer spheroids within confined environment, due to its tunable geometries and dimensions [21]. Recent works involved microfluidic devices to study spheroid mechanics via micropipette aspiration (MPA) in a high-throughput manner [22], or via compressing cellular aggregates in microchannels, to assess viscoelastic properties associated with the cell rearrangement and cell shape within the aggregate [23, 24]. Despite the valuable insights on the viscoelastic properties of tumor models gained from these studies, the relationship between the intrinsic viscoelastic characteristics of cancer spheroids and the degree of malignancy remains to be elucidated.

In this work, we establish this link by focusing on spheroids made from one non-tumorigenic epithelial cell line (MCF-10A) and two breast cancer cell lines with increasing metastatic potential(MCF-7 and MDA-MB-231). In a microfluidic chip, the spheroids are subjected to dynamic compression through a narrow constriction channel, with different level of confinement. We quantify the spheroids viscoelastic properties, by fitting the compression data to a modified Maxwell model adapted to the dynamic conditions, from which it is possible to distinguish the benign from the malignant spheroids. Then, we characterized the shape recovery of spheroids after compression during the relaxation process. Notably, comparing the fast and slow relaxation timescales of spheroids allows us to discern different degrees of malignancy, which we demonstrate to be closely associated with spheroid compactness and actin cortex arrangement.

## 2 Results

### 2.1 Microfluidic assay for dynamic compression and relaxation of breast cancer spheroids

We develop a microfluidic assay to investigate the viscoelastic response of breast cancer spheroids subjected to dynamic compression. This assay involves flowing spheroids through a narrow constriction channel connected with a relaxation chamber (see Figure 1a). The flow rate is kept constant using a syringe pump. The pump withdraws one spheroid at a time from a suspension at the micropipette loading port, compressing each spheroid through the constriction channel (referred to as the compression stage). After passing through the constriction, the spheroids are released into a wider chamber where they can relax in the absence of flow (referred to the relaxation stage). The spheroid undergoes maximum compressive strain within the constriction channel, measured by the constriction index *I*_*c*_ = (*D*_0_ − *d*_*c*_)*/D*_0_, defined by the width of the constricting channel *d*_*c*_, and the equivalent spherical diameter of the spheroid *D*_0_, such that a larger *I*_*c*_ signifies larger maximum strain. We prevent the spheroid from tumbling in the observation plane by maintaining the channel height below *D*_0_ (180 *±* 15*µ*m). We estimate the spheroid deformation during dynamic compression through the axial strain *ϵ* = (*D*(*t*) − *D*_0_)*/D*_0_, where *D*(*t*) refers to the axial dimension of the deformed spheroid over time (Figure 1b).

**Figure 1:**
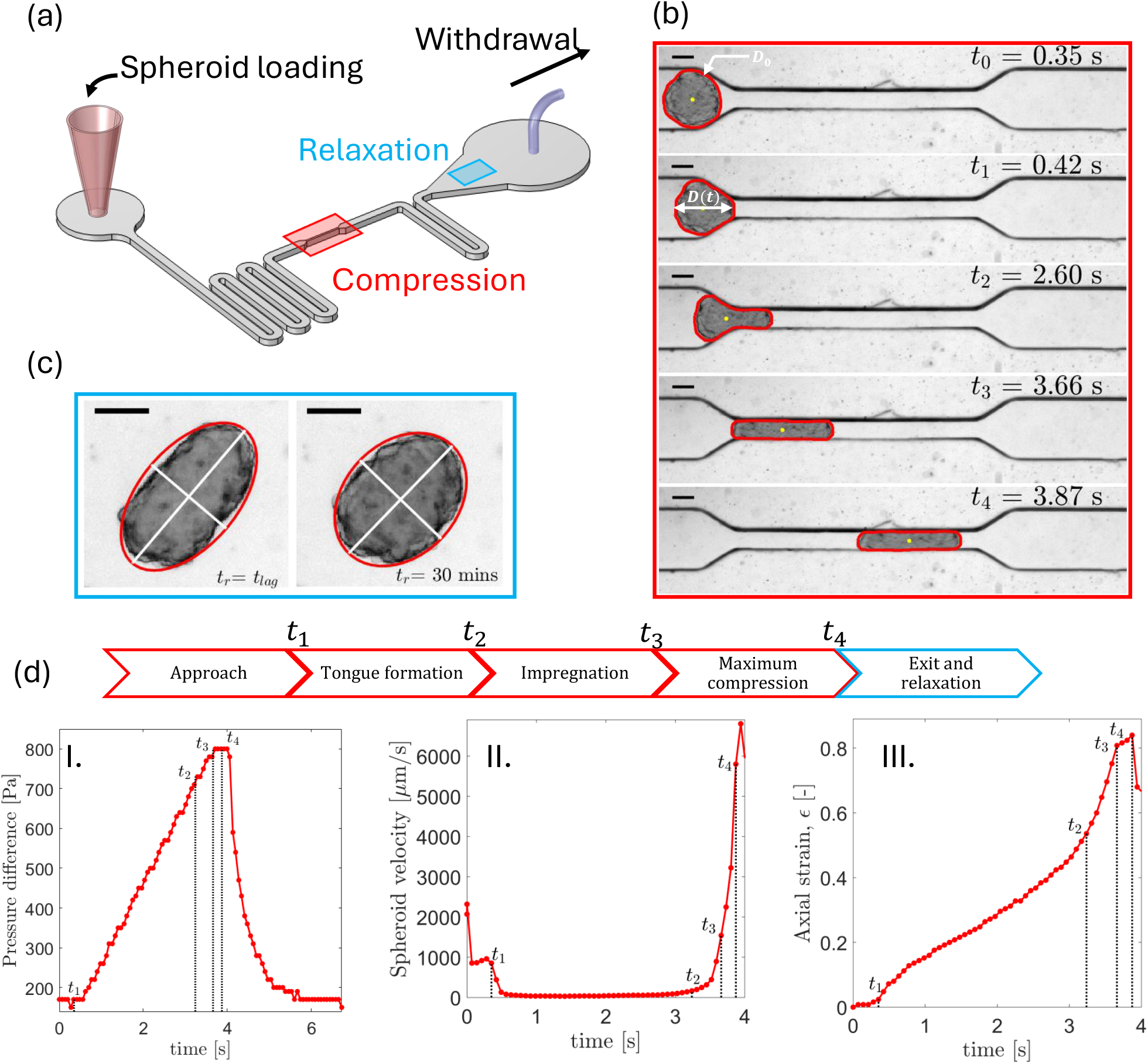
Design and working principle of the microfluidic device for viscoelastic characterization of breast spheroids with different level of malignancy. a) Schematic of the microfluidic chip for spheroids’ dynamic compression and relaxation. The rectangles identify the middle constriction (red) and the relaxation chamber (light blue). b) Showcase of MCF-7 (*I*_*c*_ = 0.63) spheroid during the compression phase, with emphasis on relevant time points. The equivalent spherical diameter *D*_0_ and the axial dimension over time *D*(*t*) are highlighted. The spheroid edge is in red and the centroid in yellow. c) Showcase of the same MCF-7 spheroid during the relaxation phase at recovery time *t*_*r*_ = *t*_*lag*_ and 30 minutes respectively. A red ellipse estimates the spheroid shape, with the major and minor axes, shown in white, tracked over time for subsequent data analysis. d) Time axis and identification of relevant time points of the deformation events and corresponding evolution of pressure difference across the channel (I), spheroids velocity (II) and axial strain (III) as function of time. Scale bar of the brightfield images: 100 *µ*m.

The dynamic compression of the spheroid can be characterized by three distinct phases: tongue aspiration (*t*_1_ *< t* ≤ *t*_2_), impregnation (*t*_2_ *< t* ≤ *t*_3_), and maximal compression (*t*_3_ *< t* ≤ *t*_4_). Figure 1d captures the dynamics of these phases for a representative MCF-7 spheroid with *I*_*c*_ = 0.63 demonstrating the corresponding time-wise variations in the (I) pressure difference across the channel (Δ*P*), (II) velocity of the spheroid (*u*_*s*_), and (III) its axial strain (*ϵ*). Immediately after clogging the entrance of the constriction channel (*t* = *t*_1_), the spheroid’s velocity drops significantly, accompanied by an increased pressure difference across the channel. This increasing Δ*P* dynamically compresses the spheroid by initiating a tongue at the leading front that is aspirated through the channel (*t > t*_1_). As soon as the spheroid’s centroid transits the constriction channel (*t* = *t*_2_), the trailing end rapidly slips into the channel, leading to complete impregnation at *t* = *t*_3_ and a sudden acceleration of the spheroid. During this phase, a rapid growth in axial strain *ϵ* takes place, reaching its maximum during the time *t*_3_ *< t* ≤ *t*_4_ as the spheroid flows inside the constriction channel. At *t* ≥ *t*_4_, the spheroid unclogs and exits from the constriction enabling fluid flow through the channel, with a consequent drop in Δ*P* and *u*_*s*_.

We monitor the relaxation of the spheroid for 30 minutes in the adjacent relaxation chamber. Unlike the constrained relaxation observed in MPA [25], the spheroids here are allowed to relax freely after exiting the constriction channel. Due to the imaging conditions and the chip design, it is not possible to capture both the constriction channel and the relaxation chamber within the same field of view. This results in a lag time *t*_lag_ ≈ 15-34 s (Figure S1) between the frames at *t*_4_, where the spheroids experience maximum compression, and the start of the relaxation monitoring (Figure 1c). We quantify the morphological relaxation of spheroid through a deformation parameter (*𝒮*_*r*_) and the circularity (*𝒞*_*r*_). Here, 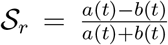 with *a* and *b* being the major and minor axes of the ellipse, respectively, that fits the 2D spheroid boundary (see Figure 1c and 3a) [26, 27, 28] (details in the Experimental Section, Data acquisition and analysis). The circularity is quantified as *𝒞*_*r*_ = (4*π ·* Area*/*Perimeter^2^).

### 2.2 Viscoelastic response as a signature of metastatic potential in spheroids under compression

To probe the link between viscoelastic properties of spheroids and their cancer malignancy, we measure the axial deformation during the dynamic compression of three different breast cancer spheroids, formed from benign cells (MCF-10A) and from low and high metastatic cells (MCF-7 and MDA-MB-231). Figure 2a shows the time evolution of axial strain (*ϵ*) for three representative spheroids of each phenotype. Notably, the spheroid from the healthy cell line (MCF-10A) exhibits the highest resistance to deformation, followed by MCF-7 and MDA-MB-231, consistent with their increasing metastatic potential (see Supporting Movie S1). The benign spheroid (MCF-10A) demonstrates a significant delay in tongue formation with *t*_1_= 68.9 s, in contrast to the two cancer spheroids (MCF-7 and MDA-MB-231) with *t*_1_ ≈ 0.07 s (as shown in Figure 2c). This results in an order of magnitude longer entry time (*t*_*E*_ = *t*_3_ − *t*_0_) (see Figure 2d), defined as the duration from when the spheroid contacts the entrance of the constriction channel (*t*_0_) until the spheroid has completely entered the middle channel at time (*t*_3_). We do not observe a statistically significant difference in *t*_*E*_ between the two malignant spheroids (MCF-7 and MDA-MB-231). Once the spheroid is fully inside the constriction channel during maximum compression, it traverses through the channel at nearly the same rate regardless of the cell type.

**Figure 2:**
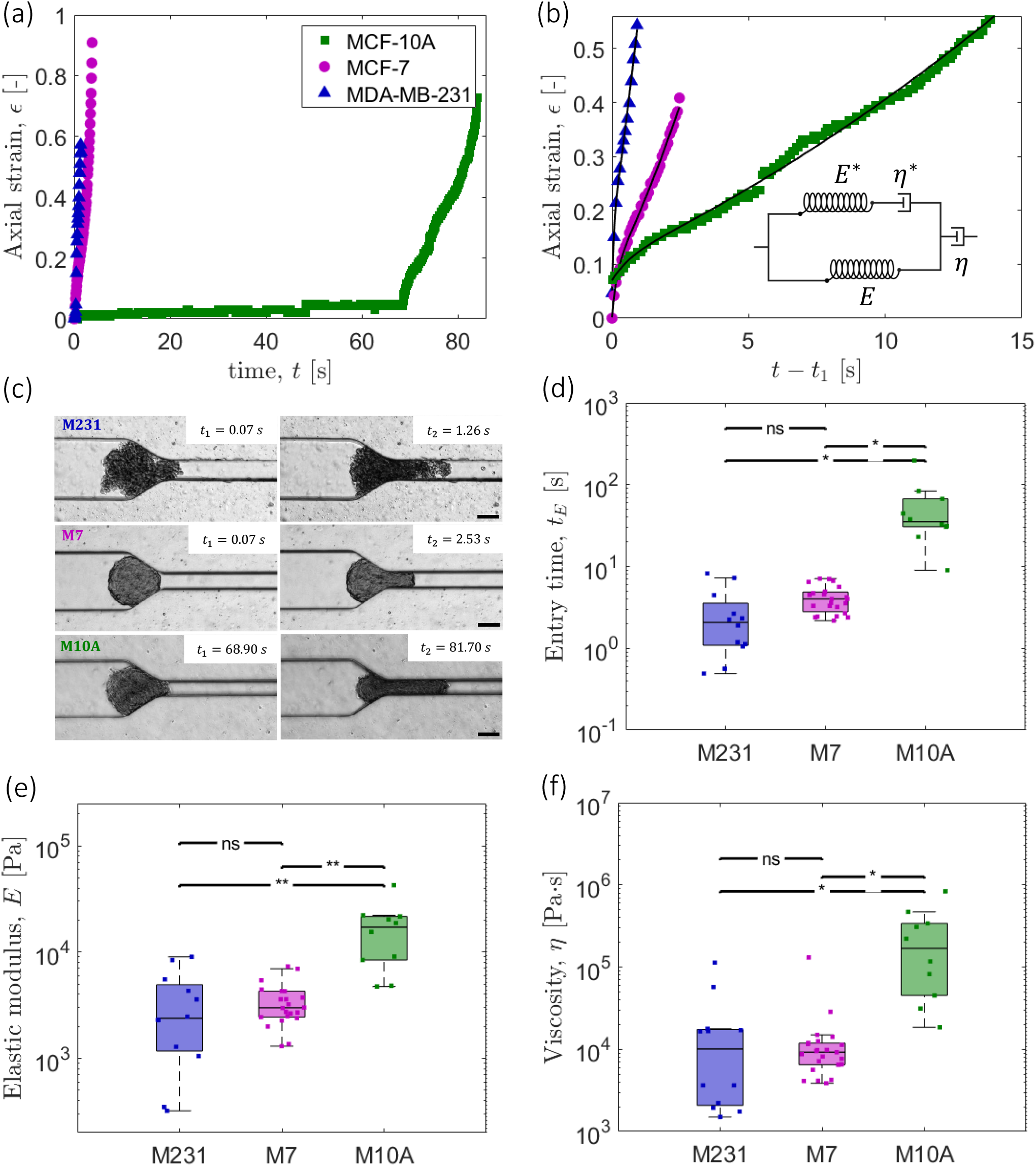
Dynamic compression of spheroid and the Dynamic Modified Maxwell model (D3M) fitting to determine spheroids viscoelastic properties. a) Time-wise evolution of axial strain during spheroid dynamic compression and deformation: The MCF-10A curve shows a delay in the rising of the axial strain, attributed to the longer time needed for tongue formation. b) The strain curves, shifted to the beginning of tongue formation *t*_1_, are fitted with the D3M, the model being illustrated in the inset. c) Brightfield images of MDA-MB-231, MCF-7 and MCF-10A spheroids in the constriction channel: snapshots taken at time *t*_1_ and *t*_2_, showing that the time for MCF-10A spheroids for tongue formation is much longer than the other two malignant spheroids. Scale bar: 100 *µ*m. d) Entry time (*t*_*E*_) of the three spheroid types for *I*_*c*_ ≥ 0.6. MDA-MB-231 and MCF-7 spheroids require a shorter time to fully enter the middle constricted channels compared to MCF-10A. e) and f) Boxplots comparing the spheroids bulk elastic moduli (E) and viscosity (*η*) for different metastatic potentials and for constriction index *I*_*c*_ ≥ 0.6.

To estimate the elastic modulus (*E*) and viscosity (*η*) of each spheroid based on the evolution of its axial strain under dynamic compression, we adapt the modified Maxwell model to account for the dynamic experimental conditions of our microfluidic assay. The modified Maxwell model was applied to MPA techniques previously to assess the viscoelastic properties of biological drops [29], and multicellular spheroids [22, 30]. The proposed model consists of four elements [31], as depicted in Figure 2b: the Kelvin-Voigt body is characterized by a spring of elasticity *E*, accounting for the bulk elasticity of the spheroid, in parallel with a spring of elasticity *E*^*^ and a dashpot of viscosity *η*^*^. The latter two describe the initial elastic jump in the strain and the local viscosity respectively. Another dashpot in series characterizes the bulk viscosity *η* of the spheroid (or aggregate). The governing empirical equation describing strain evolution in this system is given by (full derivation in SI):

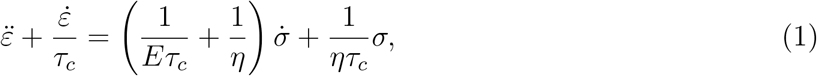

where 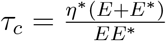 is a characteristic time for the fast elastic jump, determined by the cell-scale viscoelasticity (*E*^*^ and *η*^*^) [31]. The applied stress *σ*, due to the pressure difference across the channel, is given by 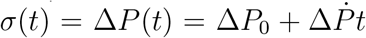. Here, 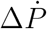 is the slope of the linear increment in the pressure (Δ*P*) against time during tongue formation (see Figure 1d (I)). The integration of Equation (1) yields the generalized equation presented in Equation (2), which we refer to as the dynamic modified Maxwell model (D3M). This model describes strain as a function of the time-dependent pressure difference (linked to the applied stress), and the viscoelastic parameters (*E* and *η*) of the spheroid:

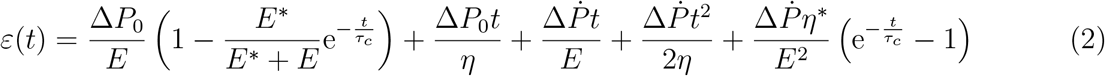

The right-hand side of equation (2) comprises five terms: the non-linear elastic jump, linear viscous and linear elastic terms, non-linear viscous term, and non-linear viscoelastic term. The analysis reported in Figure S2 shows that the last term, with *E*^2^ in the denominator, is negligible. In the limit of a constant Δ*P*, condition used in earlier studies with micropipette aspiration, we recover the modified Maxwell model equation [29]. Since D3M considers the pressure to be a linear function of time, the onset time for tongue aspiration (*t*_1_) corresponds Δ*P* = Δ*P*_0_, initial condition of the model. As illustrated in Figure 2b, this D3M model captures the strain evolution curve through the tongue aspiration phase (*t*_1_ *< t* ≤ *t*_2_) for all three spheroids. The brightfield images in Figure 2c compare the shape of each spheroid phenotype at the beginning (*t*_1_) and the end (*t*_2_) of the tongue aspiration phase. The accuracy of the D3M fit is found to be better than the modified Maxwell model (see Figure S3 and Table 1 in SI), which does not include the time-dependent linear elastic and non-linear viscous components of strain (see Equation 2). The data, as well as the model, shows a more distinct initial jump in *ϵ* for the more metastatic cell line, MDA-MB-231, which emphasizes the tendency of tumor cells to respond promptly and adapt to the deformations induced by either external forces or confinements. Immediately after the end of aspiration (when the spheroid centroid enters the constriction), the model becomes incapable to capture the sudden increase in strain, likely due to the unaccounted acceleration of the spheroid in the subsequent impregnation phase.

The viscoelastic response observed in Figure 2b is reflected in the parameters (*E* and *η*) obtained from fitting the D3M: Figures 2e-f reveal that the benign (MCF-10A) spheroids have higher elastic modulus and viscosity compared to the spheroids made of malignant cells for the constriction index ranges *I*_*c*_ ≥ 0.6. The values of elastic modulus and viscosity are in agreement with previous studies that employed MPA [22, 23], compression tests [32, 33, 34] and AFM on breast spheroids [35], which reported elastic moduli of the order of ∼ 10^2^ − 10^3^*Pa* and viscosity reaching values up to 10^5^*Pa· s*. The time-scale (*τ*_*t*_ = *η/E*) over which the spheroid transitions from the elastic to the viscous regime during compression [31] is higher for MCF-10A (median *τ*_*t*_ = 8.3 s) compared to MCF-7 (median *τ*_*c*_ = 2.9 s) and MDA-MB-231 (median *τ*_*t*_ = 4.4 s), suggesting that the benign spheroid transitions more slowly into the viscous regime than the malignant spheroids (see Figure S4).

We also compare the viscoelastic properties of MCF-7 and MCF-10A spheroids at low and high levels of compression, with *I*_*c*_ *<* 0.6 and *I*_*c*_ ≥ 0.6, respectively (see Figure S5). Notably, the spheroid viscoelasticity for MCF-10A is *I*_*c*_-dependent, showing an order of magnitude increase for both *E* and *η* as the imposed compression increases (*I*_*c*_ ≥ 0.6), in a similar trend as in single cells observed previously [36]. The effect of the constriction index for MCF-7 spheroids is minimal, with no significant difference in both *E* and *η* for lower compression levels, when *I*_*c*_ ≥ 0.6. The experiments with MDA-MB-231 cells are limited only to *I*_*c*_ ≥ 0.6 due to the difficulties in forming smaller, more compact spheroids that would fall within a lower *I*_*c*_ range.

### 2.3 Relaxation response and shape recovery of spheroids with different metastatic potentials

To test if the relaxation behavior of the spheroids also bears signatures of metastatic potential, we analyze their subsequent shape recovery in the absence of flow. Figure 3(a) shows representative images of three spheroids at the first (*t* = *t*_lag_) and last (*t*=30 min) captured frames in the relaxation chamber. Figure 3b and 3c illustrate the typical relaxation of MDA-MB-231, MCF-7 and MCF-10A spheroids through the time-wise evolution in dimensionless deformation parameter (*S*_*r*_) and circularity (*𝒞*_*r*_). For both curves, the starting points correspond to the values of 𝒮_*r*_ and *𝒞*_*r*_ at the moment of maximum compression at *t* = *t*_4_. This allows for the retrieval of the spheroid status immediately preceding the start of relaxation despite the lag time. After a fast rebound (see inset of Figure 3b), all three spheroids relax slowly and asymptotically toward a plateau (see Supporting Movie S2). In contrast to healthy MCF-10A spheroid, the malignant MCF-7 spheroid continues to relax and does not retrieve its original deformation (𝒮_*r*0_) until the end of the experiment (≈30 min). This results in a residual deformation (𝒮_*r*∞_ *> 𝒮* _*r*0_) for MCF-7, associated with long time viscous effects. Interestingly, the MDA-MB-231 spheroid shows a different relaxation behavior compared to the other two cell-types. The MDA-MB-231 (with mesenchymal features) spheroids relaxes through rearrangement of constituent cells, such that the final deformation parameter (𝒮_*r*∞_) fluctuates significantly among cases and even reduces below its pre-compression value 𝒮_*r*0_ in the example shown in Figure 3b. To quantify the fast and slow relaxation and the subsequent residual deformation of breast spheroids, we employ a double exponential model (DEM) fit to the measured temporal deformation parameter (𝒮_*r*_), given by:

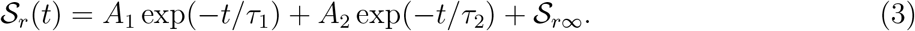

**Figure 3:**
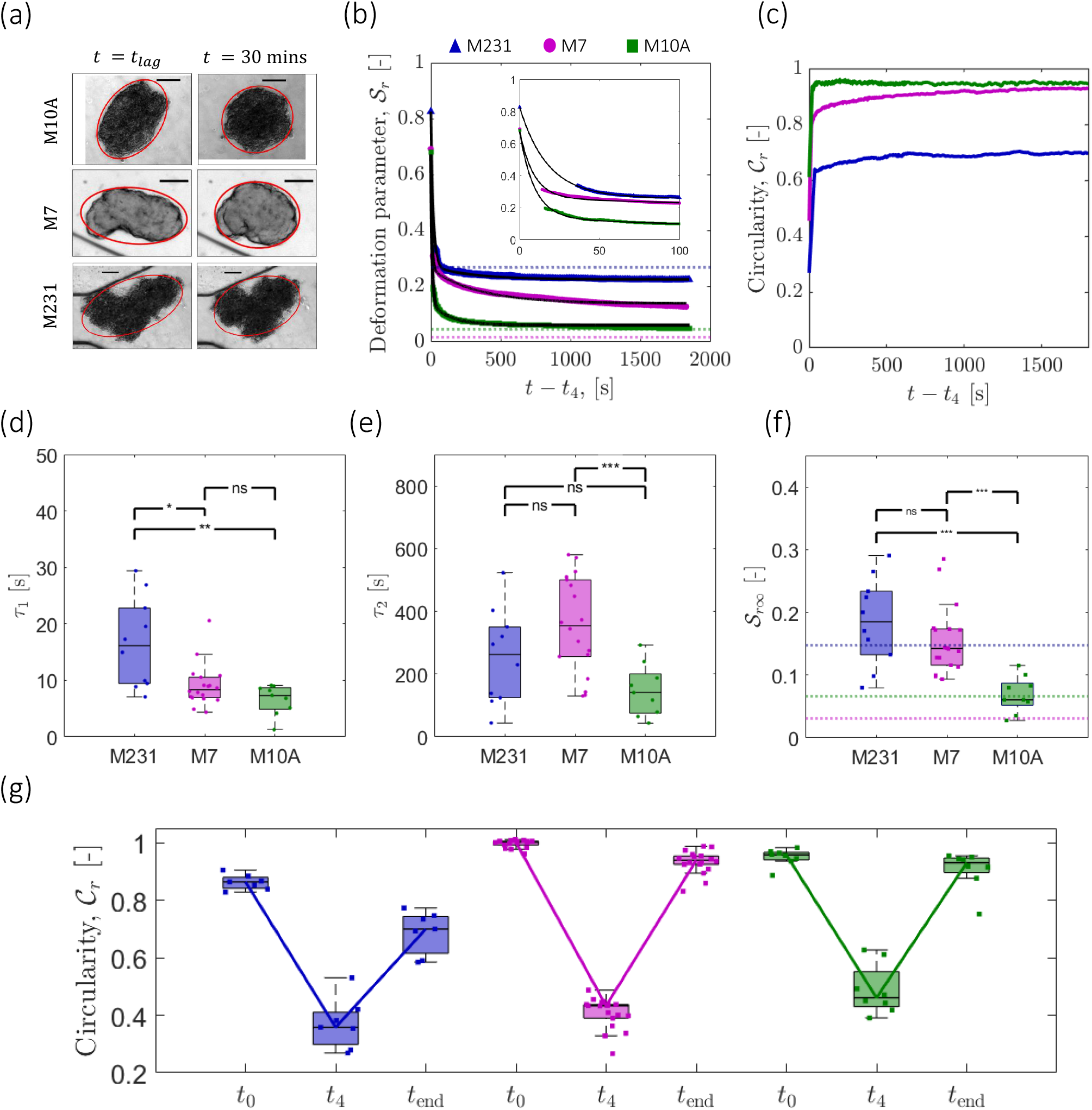
Shape relaxation and recovery of spheroids with different metastatic potential after the dynamic compression (*I*_*c*_ ≥ 0.6). a) Brightfield images of MDA-MB-231, MCF-7 and MCF-10A spheroids in the relaxation phase: images taken at t = lag time and after 30 minutes from the time of maximum compression *t*_4_. b) Deformation parameter 𝒮_*r*_ over time and the double exponential model (DEM) fitting. The starting point in each curve is shifted to *t*_4_ and refers to the 𝒮_*r*_ value at maximum compression. The dotted lines refer to the deformation parameter of each spheroid prior to compression 𝒮_*r*0_. The inset shows the fitting of the first 100 seconds. c) Circularity *𝒞*_*r*_ curve over time during the relaxation phase. MCF-10A and MCF-7 spheroids recover the circular shape, whereas MDA-MB-231 spheroids maintain an irregular shape after deformation. d) Short (*τ*_1_) e) long (*τ*_2_) timescales of relaxation and f) residual deformation 𝒮_*r*∞_ based on the DEM fits for spheroids of *I*_*c*_ ≥ 0.6. The dotted lines in 3f indicate the median values of the deformation parameter prior to compression 𝒮_*r*0_. g) Boxplots of circularity *𝒞*_*r*_ before compression (*t*_0_), at the maximum compression (*t*_4_) and after 30 minutes of relaxation (*t*_*end*_) for the three spheroid type. The highly metastatic spheroids do not recover their initial circularity values, differently from the less metastatic MCF-7 spheroids and the benign MCF-10A spheroids.

The DEM characterizes the multiscale relaxation process via the two time-scales: *τ*_1_ (short) and *τ*_2_ (long), and the residual deformation (𝒮_*r*∞_) as fitting parameters. Herein, *τ*_1_ corresponds to the immediate rebound of the spheroid after exiting from the constriction channel, and *τ*_2_ to the subsequent long-term local rearrangements, likely associated with inter-cellular interactions. Both breast cancer single cells and other non-cancerous compressed multicellular aggregates are known to exhibit such double exponential behavior as observed in earlier studies [37, 38, 32, 39, 40], based on other techniques like tissue surface tensiometry and AFM. For the spheroids used in our study, the relaxation curves are better captured by the DEM compared to some other fitting models, such as a single exponential and power-law fittings [41, 42], as shown in Figure S6.

All three types of spheroids show an early-time fast elastic rebound with *τ*_1_ of the order of ≈10 seconds, indicating an immediate elastic response after exiting the constriction. We find this timescale to be the shortest for the benign spheroid (MCF-10A), followed by progressively longer *τ*_1_ corresponding to malignant spheroids (MCF-7 and MDA-MB-231), as shown in Figure 3d. The statistical difference in *τ*_1_ among the three spheroid is significant only when comparing the benign (MCF-10A) and low-metastatic (MCF-7) spheroids with the highly metastatic (MDA-MB-231) spheroids. Notably, both MCF-7 and MCF-10A can be distinguished from highly metastatic MDA-MB-231 spheroids in terms of greater compactness, quantified by their high circularity values (*𝒞*_*r*_ ≈ 1) prior to compression (Figure 3g). To examine this further, we visualize the F-actin distribution within each spheroid prior to compression (Figure 4a-b). We observe that both MCF-10A and MCF-7 form compact spheroids distinguished by a peripheral actin rim, which is more pronounced in the benign MCF-10A spheroids. Based on this result, we propose that the fast relaxation (i.e. *τ*_1_) is primarily influenced by surface elasticity resulting from the peripheral distribution of actin fibers, rather than solely by the bulk elasticity. This is supported by the significant difference in bulk elastic moduli between MCF-10A and MCF-7. By contrast, MDA-MB-231, which exhibit lower circularity (*𝒞*_*r*_ ≈ 0.85) and lack a visible F-actin cortex, displays prolonged initial relaxation and greater variability in the *τ*_1_ values. This suggests that the absence of a structured actin cortex contributes to a more heterogeneous and extended relaxation response in the MDA-MB-231 spheroid (see inset in Figure 3b).

**Figure 4:**
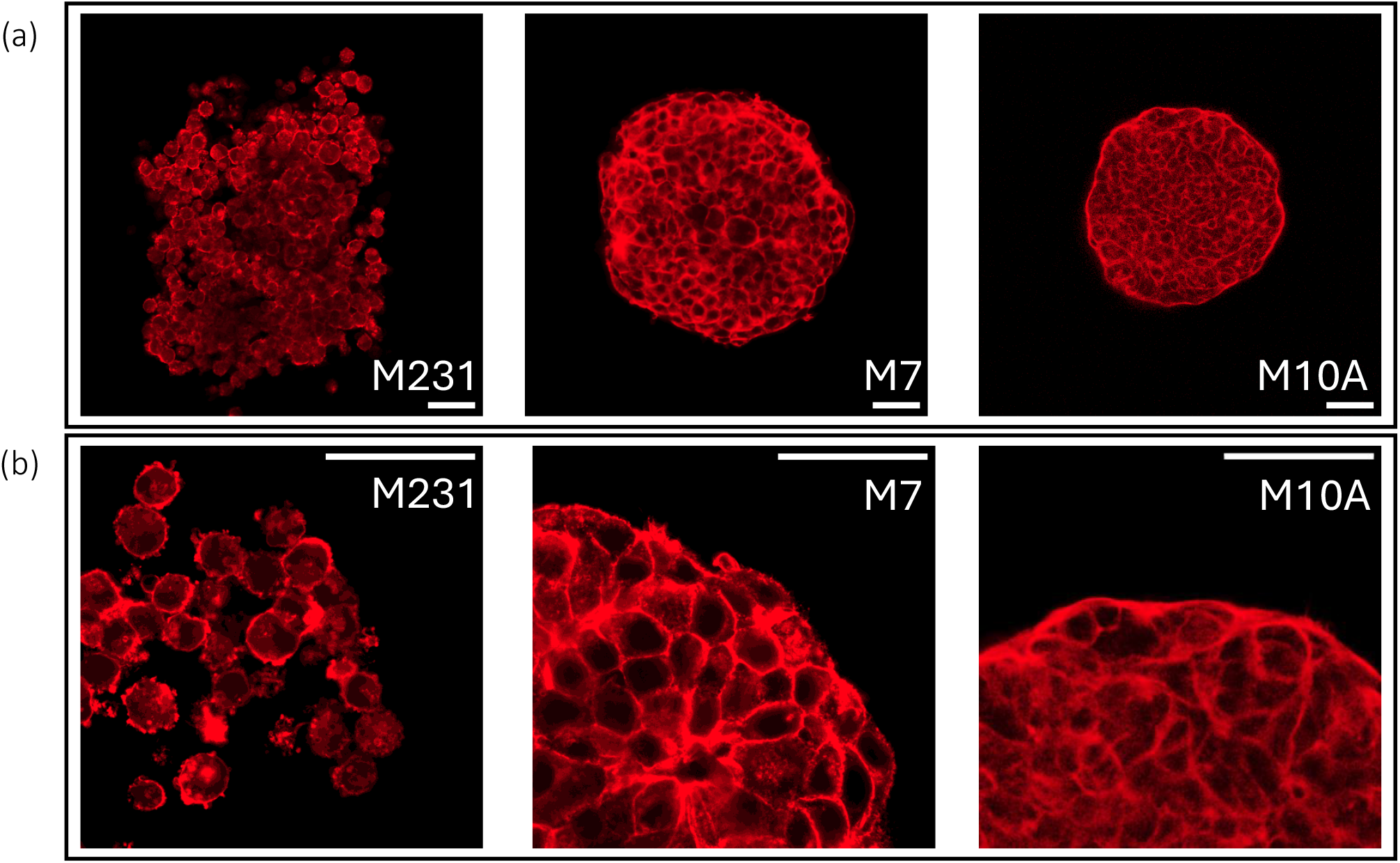
F-actin immunostaining in breast cancer spheroids of different malignancy. (a) Confocal images (at 20x) of F-actin (at the equitorial plane) in MDA-MB-231, MCF-7 and MCF-10A breast spheroids (from left to right): MCF-10A spheroids display smooth edges with a prominent collective actin rim, MCF-7 spheroids show similar but less pronounced actin rim, while MDA-MB-231 spheroids exhibit fragmented edges lacking a collective actin rim. (b) Same spheroids at higher magnification (63x): MCF-10A cells at the boundary display stretched morphology with compact and smooth actin organization. MCF-7 spheroid edges are similarly compact, but have slightly rougher contours. In contrast, MDA-MB-231 spheroids feature non-interconnected cells with irregular F-actin distribution. Scale bar is 50 *µ*m.

The subsequent slow relaxation is characterized by the longer timescale *τ*_2_, governed by the bulk elasticity (*E*). Benign spheroids (MCF-10A) exhibit the shortest relaxation time (with *τ*_2_ *<* 141 s), indicative of their higher elasticity and compact structure, enabling quick and uniform relaxation. The two malignant cell lines (MCF-7 and MDA-MB-231) display similar slow relaxation times, but with higher *τ*_2_ compared to MCF-10A. These late-time relaxation timescales are consistent with the *E* values derived from dynamic compression and deformation, with lower elasticity for the malignant spheroids (MCF-7 and MDA-MB-231) compared to MCF-10A spheroids. The median value of *τ*_2_ for MDA-MB-231 being smaller than more MCF-7 is reflective of a more compliant and flexible spheroid, with cells at the periphery that are more free to move and rearrange, while MCF-7 spheroids, despite being less metastatic, maintain their deformed state for a longer time, but are still capable to recover most of the spherical initial shape (up to 93.7%), as shown by the circularity plot in Figure 3g.

At the end of 30 minutes of relaxation, all benign MCF-10A spheroids nearly regain their original deformation 𝒮_*r*0_, such that 𝒮_*r*∞_ → 𝒮_*r*0_ (Figure 3f). They also achieve a final circularity value nearly identical to their initial shape before compression (retrieving 97% of original circularity), as depicted in Figure 3g. By contrast, the malignant MCF-7 cell lines display much greater resistance to recovery due to long time viscous effects with significant residual deformation (𝒮_*r*∞_) at the end of our experimental time. The residual deformation of MDA-MB-231 (𝒮_*r*∞_) is found to be case specific due to cell-cell rearrangements, which results in a broad range of 𝒮_*r*∞_ values both above and below their original deformation parameter (𝒮_*r*0_). The two malignant spheroids also differ significantly in their morphological recovery: MCF-7 spheroids regain a spherical shape, while MDA-MB-231 spheroids display fragmented edges, with cells moving in different directions (Supporting Movie S2), leading to an overall irregular spheroid shape.

## 3 Discussion and Conclusion

In this work, we investigate the relationship between the viscoelastic properties of breast spheroids and their metastatic potential through dynamic compression and relaxation experiments. We use a microfluidic chip to systematically analyze spheroid deformation and subsequent relaxation, allowing us to obtain the bulk elastic modulus (*E*) and viscosity (*η*) through a viscoelastic model tailored to the experimental conditions.

Benign spheroids display higher *E* and *η* compared to the low and high metastatic counterparts (MCF-7, MDA-MB-231). These findings on breast cancer spheroids align with previous studies on single breast cancer cells [43, 36, 44, 45] and on spheroids of different tissues, e.g. originating from bladder cancer [46]. Notably, among the three cell lines, only MCF-10A shows significant sensitivity in both *E* and *η* to the imposed maximum strain, with higher values corresponding to increased level of maximum strain. This behavior, also observed on single MCF-10A cells previously [36], indicates that the apparent viscoelasticity of a spheroid is not only imposed strain-dependent, but also is specific to spheroid metastatic potential.

Additionally, the variation in spheroid entry time into the constriction relative to metastatic potential corroborates earlier findings on single breast cancer cells in a similar constriction channel [9]. The higher entry time for benign MCF-10A cells, compared to MCF-7 and MDA-MB-231, was attributed to their greater stiffness, and correlates with both bulk elasticity and viscosity (Figure S7). Overall, these studies suggests that malignant breast cancer spheroids, similar to their constituent single cells, are more compliant to deformation compared to healthy spheroids.

During the relaxation phase we identify two distinct time scales for all three spheroid types. Despite the similarity in the bulk properties of both two malignant spheroids (MCF-7 and MDA-MB-231) during compression, we observe distinct behaviors in their relaxation. The early-time fast relaxation, characterized by smaller *τ*_1_ values, is attributed to surface elasticity. Both MCF-7 and MCF-10A spheroids, with their distinct peripheral actin rims and spherical shape, exhibit a faster initial rebound compared to MDA-MB-231 spheroids. Immunostaining of F-actin fibers reveals that this peripheral actin rim is present in MCF-10A and MCF-7 spheroids, but is absent in MDA-MB-231 spheroids, which exhibit a highly irregular shape. This indicates that the structural arrangement of actin fibers significantly influences the early-time relaxation dynamics making the less malignant MCF-7 behave more like the benign MCF-10A spheroid than the highly malignant MDA-MB-231 spheroid. We quantify this surface characteristic through spheroid circularity, which is much closer to 1 for MCF-10A (*𝒞*_*r*_ ≈ 0.96) and MCF-7 (*𝒞*_*r*_ ≈ 1) than MDA-MB-231 (*𝒞*_*r*_ ≈ 0.86). The late-time slow relaxation is correlated with the bulk elasticity (*E*) values with a shorter *τ*_2_ for the benign MCF-10A than the malignant MCF-7. Notably, we do not observe a dominant viscous effect within the experimental time frame of relaxation (30 minutes), especially for MCF-7 and MCF-10A. Despite MCF-10A spheroids having the highest viscosity among the three types, a longer viscous relaxation timescale is expected, but is not observed in our experiments.

Although MDA-MB-231 show a *τ*_2_ similar to MCF-7, aligning with their respective elastic moduli, we believe that the relaxation of the former is more complex and is likely influenced by the rearrangement of individual cells. The non-uniform actin distribution in MDA-MB-231 spheroid, seen in Figure 4 (also reported earlier in [47, 48]), limits its ability to recover a morphologically compact, circular shape, unlike MCF-7 (see Figures 3g and 4). Cells at the periphery of MDA-MB-231 spheroids exhibit increased mobility (Supporting Movie S2), enhancing the potential for cell-cell rearrangements. Furthermore, the lack of E-cadherin expression, which is crucial for cell-cell adhesion, in highly metastatic MDA-MB-231 cells [49, 50] contributes to the spheroid’s loosening during dynamic compression [51]. In a separate experiment, an MDA-MB-231 spheroid subjected to high *I*_*c*_ = 0.87 fragment during dynamic compression, with individual cells disseminating from the spheroid (see Movie S3). This confirms that spheroids with high metastatic potential (with mesenchymal features) are more prone to deforming irreversibly and breaking, disseminating invasive tumor cells under external physical forces (such as compression), due to unstable cell-cell contact and weak cortical contractility [47].

To summarize, we compare the viscoelastic properties of benign spheroids with malignant spheroids (with low and high metastatic potentials). Notably, the benign spheroids exhibit the highest bulk elasticity, viscosity, and resistance to deformation, in contrast to the two malignant spheroids. We find that both low and high metastatic spheroids have similar apparent viscoelastic properties; however, they differ significantly during the relaxation phase. Important extension of this work could be the investigation of the mechanical properties of patient derived biopsy samples (from patients diagnosed by breast cancer) and link them to different stages of metastasis. Our microfluidics assay is a suitable platform for studying the mechanics of a wide range of solid tissues. It provides mechanistic insights into physiological processes, such as cellular remodeling within invasive tumors, and identifies unique ”mechanical biomarkers.” These biomarkers can be applied in developing therapeutic approaches that target the dissemination of primary solid tumors [5, 6].

## 4 Experimental Section

### Cell culture and spheroid formation

Human mammary MCF-10A cells (ATCC CRL-10317) were cultured in DMEM/F12 1:1 medium (Gibco) supplemented with 5% horse serum (Gibco), 0.5*µ*g mL^−1^ hydrocortisone (Sigma), 20 ng mL^−1^ human epidermal growth factor (hEGF) (Peprotech), 100 ng mL^−1^ cholera toxin (Sigma), 10 g mL^−1^ insulin (Human Recombinant Zinc, Gibco) and 1% Penicillin–Streptomycin 100× solution (VWR Life Science). Human mammary MCF-7 mCherry cells and MDA-MB-231 LifeAct GFP (kindly provided by Peter ten Dijke’s lab) were cultured in DMEM medium (Gibco) supplemented with 5% fetal bovine serum (Gibco) and 1% antibiotic-antimycotic solution (Gibco). All cells were incubated at 37°C with 5% CO_2_ and subcultured at least twice a week. Cells were regularly tested for absence of mycoplasma. Spheroids were formed in a commercially available Corning^®^ Elplasia ^®^ 96-well plate or Sphericallate 5D ^®^ (Kugelmeiers) designed for efficient spheroid production. These plates have a round-bottom shape and a specialized Ultra-Low Attachment (ULA) surface that prevents cells from attaching to the plate and encourages cell-to-cell adhesion. The size of the spheroids depends on factors such as the initial seeding density and the duration of the culture (related to proliferation rate of each cell line). The seeding density was tuned to obtain a spheroid diameter of 200-220 *µ*m after 2 days of culture in the case of MCF-7 and MCF-10A. To promote cell adhesion and spheroid formation for MDA-MB-231 cells, the media was supplemented with methylcellulose (Merck Sigma) in a ratio 1:4 to promote spheroid compactness [52]. The spheroids were harvested after 5 days.

### Spheroids Fixation, Permeabilization, Immunostaining and Imaging

Spheroids were fixed with 4% paraformaldehyde for 30 min at room temperature (RT) and then permeabilized using 0.1% Triton X-100 (diluted in PBS) at RT for 5 min. For immunostaining, cells were incubated with Phalloidin iFluor 647 Reagent (ab176759) diluted 1:100 in 1% BSA for 90 min at RT. In between all steps, the samples were washed multiple times using PBS. Samples were kept at 4°C, protected from light. Soon after the staining procedure, the spheroids were placed in uncoated chamber wells (ibidi) for visualization of the F-actin organization. Images were acquired on a confocal microscope (LSM 980 with Airyscan 2, Zeiss) equipped with Plan Apochromat 20x/0.8 M27 air objective (to image the whole spheroid) and Plan Apochromat 63x/1.40 Oil DIC M27 objective for the spheroids’ edges (*λ*_*ex*_ = 653 nm and *λ*_*em*_ = 668 nm). Z-stacks were acquired with a step of 5*µ*m and the equitorial planes were chosen for the visualization of the whole spheroids and their edges. Images post-processing was performed on ImageJ (v1.53t, National Institute of Health, USA).

### Microfluidic chip design and experimental setup

A 4-inch silicon wafer was used as the base material for the chip, and the fabrication process was carried out in a cleanroom facility (Kavli Nanolab Delft) via a photo-lithography process using the µMLA Laser Writer (Heidelberg Instruments). A detailed procedure of the the wafer fabrication can be found in the Supporting Information. Using the master mold as a template, the microfluidic chips were then produced using the soft lithography technique with polydimethylsiloxane (PDMS) as the main material. The day before each experiment, the chips were coated with 1 % bovine serum albumin (BSA, Sigma Aldrich) in DPBS buffer solution to reduce friction of the spheroids with the channel walls. The chips were kept overnight at 37°C and with sufficient humidity. The day of the experiment the chips were flushed with DPBS to remove BSA extra residues and eventual dirt. The three serpentines positioned at the inlet allow for the stabilization of the flow when the spheroid enters the chip, whereas the last serpentine is strategically utilized to impede the spheroid’s exit, ensuring its prolonged residence in the relaxation chamber. The height of the chip (180±15 *µ*m) is uniform along the whole design. To prevent the tumbling of the spheroids, we selected spheroids with a higher diameter than the height of the chip. This allows for a pre-compression of the spheroid in the z-axis, enabling subsequent observations and measurements to be conducted within the recording x-y plane. This also implies the clogging of the flow when the spheroids enter the constriction, excluding possibilities of shear flow between the spheroids and the channel walls. The microfluidic chips have been fabricated with different constriction widths (60-65 *µ*m, 84-94 *µ*m and 100-110 *µ*m). The experimental setup consists of a 200 *µ*L pipette tip plugged into the inlet for the spheroids loading, while at the outlet a tube is connected to an In-Line pressure sensor, S version (Fluigent) for a continuous measurement of pressure. Additionally, a syringe pump (Harvard Apparatus, Pump 11 Pico Plus Elite) operating in withdrawal mode (4 *µ*L min^−1^) is used to ensure constant fluid flow and enable the spheroids to pass through the constriction in a controlled manner. When the spheroid exits the constriction the withdrawal flow rate is immediately set to zero, allowing to keep the spheroid within the relaxation chamber. The spheroids are individually loaded into the pipette tip at the inlet, together with the cell culture media, one at a time, to prevent flow and pressure instability given by the traverse of two or more spheroids in the microfluidic chip. Experiments that had more than one spheroid flowing in the chip were not considered in the analysis. All experiments were conducted at 37°C through a Heating Insert (ibidi).

### Data acquisition and analysis

Brightfield images of the spheroids entering the constricted channels were captured on an inverted fluorescence microscope (Zeiss Axio-Observer) with camera streaming (for the compression stage) using a 5×/NA 0.16 air objective and Zeiss Axio-Observer 0.63x digital camera with a resolution of 2048×2048 *µ*m^2^. Once the spheroid exited the constriction, the syringe pump was paused to allow the spheroid to reside in the chamber. For the relaxation phase, images were taken at 2 fps for the first 10 minutes, and successively at 0.1 fps for the remaining 20 minutes, for a total of 30 minutes of relaxation recording in the same imaging conditions as during the compression. To analyze the dynamic compression data, an image without the spheroid was subtracted from the image containing the spheroid to remove all background noise and identify the spheroid contour (edges in red in Figure 1b). The image was then converted to binary format, and morphological closing was applied to remove the rough edges from the boundary of the spheroid. To determine the spheroid axial dimensions prior to (D_0_) and during compression (D(t)), a MATLAB function that detects the minimum and maximum pixel values in the horizontal direction was used. The velocity (*u*) of the spheroid was calculated by first identifying the centroid (using regionprops function in MATLAB) and then averaging the displacements of the centroid for consecutive images according to the Δt from one image to the other. For the relaxation, given the variable orientation of the spheroid in the relaxation chamber, it was not possible to implement the same function used for the dynamic compression to detect the length of the deformed spheroid. Therefore, an ellipse fitting was used and its major and minor axes (white lines in Figure 1c) were detected to calculate the deformation parameter 𝒮_∇_.

### Pressure characterization and D3M model

The spheroid entering the constriction clogged the fluid flow, causing the volumetric flow rate in the channel to go to zero. However, the withdrawal force applied by the syringe pump remained constant, causing an increase in the pressure in the constriction, detected by the pressure sensor located at the outlet. The sensor recorded a linear pressure increase, meaning that the stress experienced by the spheroid linearly increased during the compression phase (Figure 2a). The deformation of the spheroid in the main constriction is fitted by the Dynamic Modified Maxwell Model (D3M), up to the inflex point of the strain curve, corresponding to the time *t*_2_. The model considers the pressure as an affine line, rather than a constant value as the Modified Maxwell Model (MMM), used for conventional MPA experiments. The two models share the same schematic, but the governing equations are different because of the time dependence of the pressure.

The general governing equation is given by (for full derivation refer to Supporting Information):

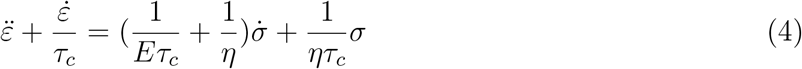

with 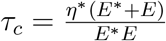

The initial conditions for the stress and strain are the following:

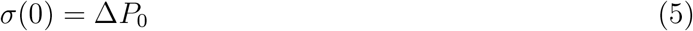

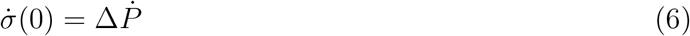

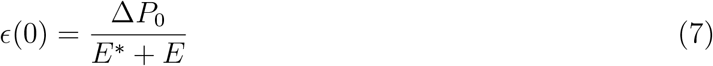

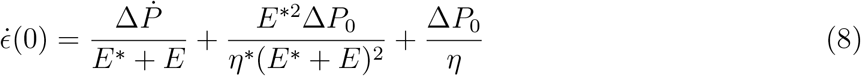

The resolution of the differential equation gives us:

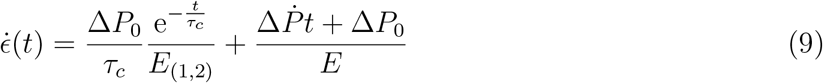

with 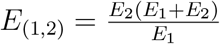

### Statistical analysis

All statistical analysis was performed using Microsoft Excel (Microsoft Corporation, USA) and MATLAB. The statistical significant differences between the experimental groups were determined by Student t-test using the function t-test: two samples with unequal variance and p values below 0.05 were considered to be significant. We categorize statistical differences as following; p *<* 0.001 (***), p *<* 0.01 (**) and p *<* 0.05 (*).

## Supporting Information

Supporting Information is available from the Wiley Online Library or from the author.

## Author Contributions

M. T., A. D. B. and P. E. B. conceived the ideas and designed the experiments. M. T., E. T., S. D. carried out the experiments and collected the data. M. T. and A. D. B. performed data and image analysis. A. D. B. developed MATLAB codes for image analysis. E.T., A.D.B. and M.T. developed the model. M. T. wrote the paper and A. D. B., V. G. and P. E. B. edited it.

## Acknowledgements

The authors thank Peter ten Dijke (from Leiden University Medical Center) for kindly providing MCF-7 mCherry and MDA-MB-231 LifeAct GFP cells. M.T. and P. E. B. gratefully acknowledge funding from the European Research Council (ERC) under the European Union’s Horizon 2020 research and innovation program (grant agreement no. 819424). A. D. B. gratefully acknowledges funding from MSCA Postdoctoral Fellowships 2022 Project ID: 101111247. E.T. was supported by an Erasmus+ Traineeship.

## Data Availability Statement

The data that support the findings of this study are available from the corresponding author upon reasonable request.

## Supporting information

### Details about microfluidic chip fabrication

The master wafer was fabricated by a standard photo-lithography technique using a µMLA laserwriter (Heidelberg Instruments) on a 4-inch silicon wafer at the Kavli Nanolab Delft.

We first spin-coated SU-8 2100 (Kayaku Advanced Materials) negative photoresist in two steps:

1. 500 rpm for 10 seconds with an acceleration of 100 rpm/s
2. 1850 rpm for 30 seconds with an acceleration of 300 rpm/s.

The silicon-wafer with SU-8 was soft-baked at 65 °C for 5 minutes then at 95 °C for 30 minutes. The wafer was then inserted in the laser writer holder for printing using a 365mn laser source. Details of the AutoCAD design can be found here (https://github.com/margheritatavasso/Microfluidicdesign_compression_relaxation-).

After laser printing, the wafer was post baked at 65 °C for 5 minutes, followed by 95 °C for 12 minutes and developed in SU-8 photo-resist developer (mr-Dev 600, Micro resist technology or PGMEA, Sigma Aldrich). After development, a hard bake at 150°C for 5 minutes ensured better defined patterns. We optimized the photolithography procedure to achieve a final channel height of 180 ± 15 µm. The height of the channels was determined by a Dektak profilometer.

The master wafer was coated with trichloro(1H,1H,2H,2H-perfluorooctyl)silane to create a hydrophobic surface for easy demoulding. Polydimethylsiloxane (PDMS, Sylgard 184) based microfluidic chips were prepared using the mixture of PDMS and curing agent with a ratio of 10:1. Individual chips were then cut out and inlets/outlets were punched in the PDMS slab for tube fitting (PTFE, inner diameter: 0.3mm, outer diameter: 1.6mm). Final bonding is performed by plasma cleaning (Harrick Plasma) of the PDMS slab and a glass coverslip (#1.5) for two 2 minutes 20 seconds to facilitate final bonding. Finally, the assembled device was kept in the oven at 70 °C for at least 20 minutes to facilitate bonding strength. Afterwards, the microfluidic devices can be stored indefinitely.

### Derivation of the Dynamic Modified Maxwell Model

The schematic of the model is the following:

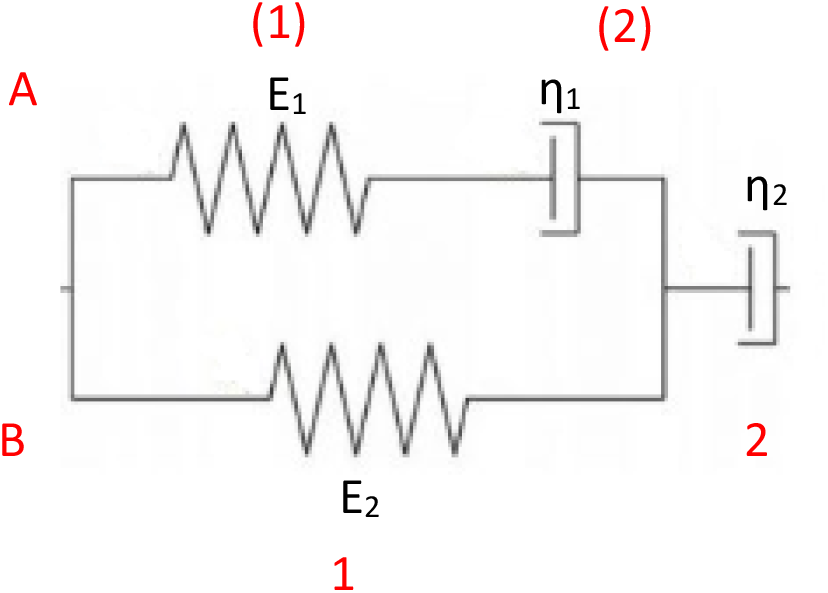

For the springs in the model Hooke’s law applies:

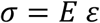

The dashpots follow Newton’s law for Newtonian fluids, where

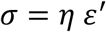

Based on these, we have the following equations

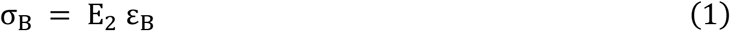

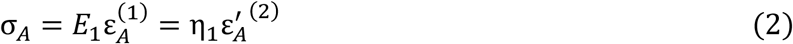

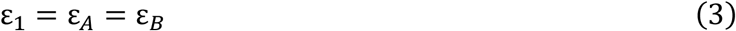

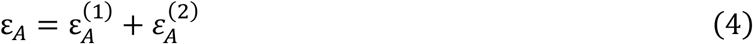

From Equation (3) and Equation (2) we can respectively write

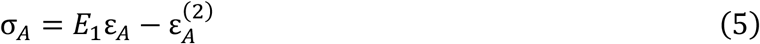

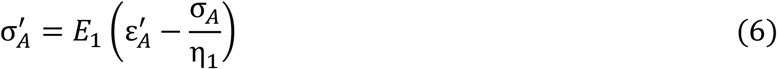

With σ_1_ = σ_*A*_ = σ_*B*_ (7) and therefore 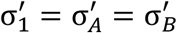 we can write that

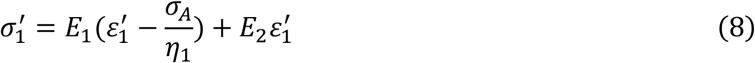

By implementing Eq (7) in the first term

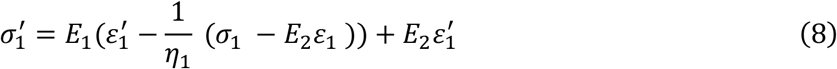

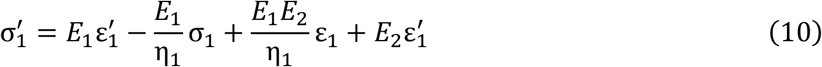

Given that σ = σ_1_ = σ_2_ and ε = ε_1_ + ε_2_ we have that ε_1_ = ε − ε_2_. We can rewrite Eq (10) as

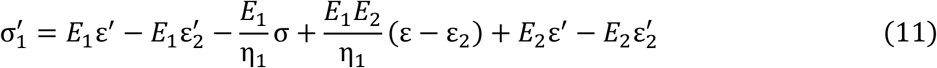

Considering the second derivative of the stress *σ*_1_ (which is 0) and rewriting the equation with the strain terms on the left side and the stress on the right side of the equation, we obtain:

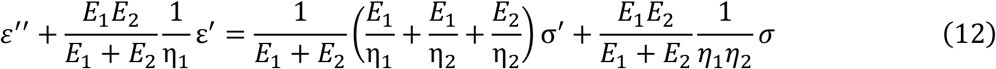

This equation can be rewritten as the following:

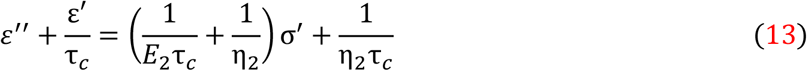

with 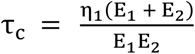

This is the constitutive equation of the model that will be solved with the appropriate initial conditions.

### Solving the differential equation with initial conditions

For the stress on the spheroid, the following expression is used: σ(*t*) = Δ*P*(*t*) = *at* + Δ*P*_0_

To simplify we write σ = *at* + *b* and therefore *σ*^′^ = *a*.

By substituting these two formulas in Eq 13 we obtain:

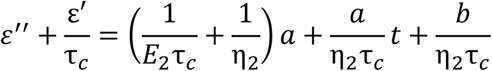

The following initial conditions are used to solve this differential equation:

1. 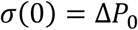
2. 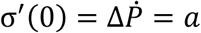
3. 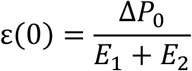
4. 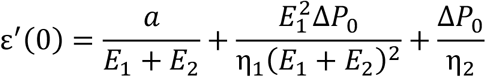

The last initial condition is obtained by solving 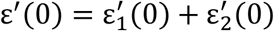with 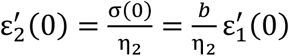 and 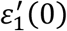 is taken by rewriting Eq(10) and substituting first two initial conditions.

The homogeneous solution for the strain derivative is 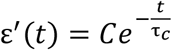

From Eq(13) the particular solution is:

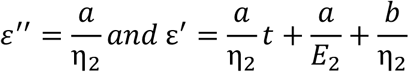

When we combine the homogeneous and particular solution we obtain:

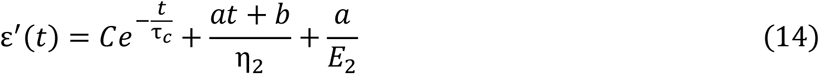

We substitute first the 4. Initial condition, this gives us the constant C from the homogeneous solution:

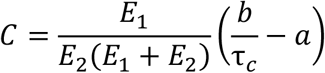

We define 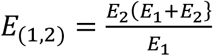 and substitute the value of the constant C in Eq(14):

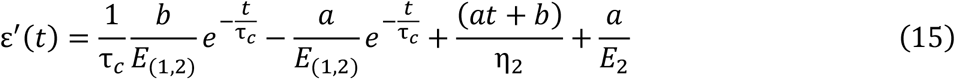

We integrate Eq(15) into the following:

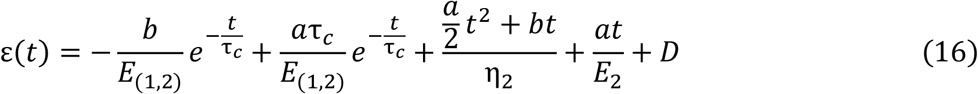

We implement the 3. Initial condition that gives us the value of the constant D:

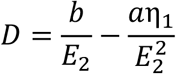

By substituting the value of D and rewriting 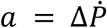, *E*_1_ = *E*^*^, *E*_2_ = *E, η*_1_ = *η* * *and η*_2_ = *η* we obtain:

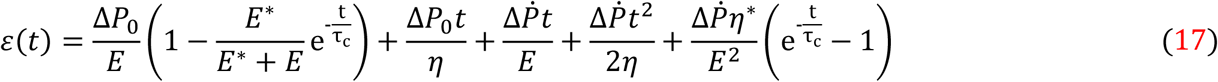

Which is the final equation of the Dynamic Modified Maxwell Model.

## Supplementary Figures

**Figure S1.**
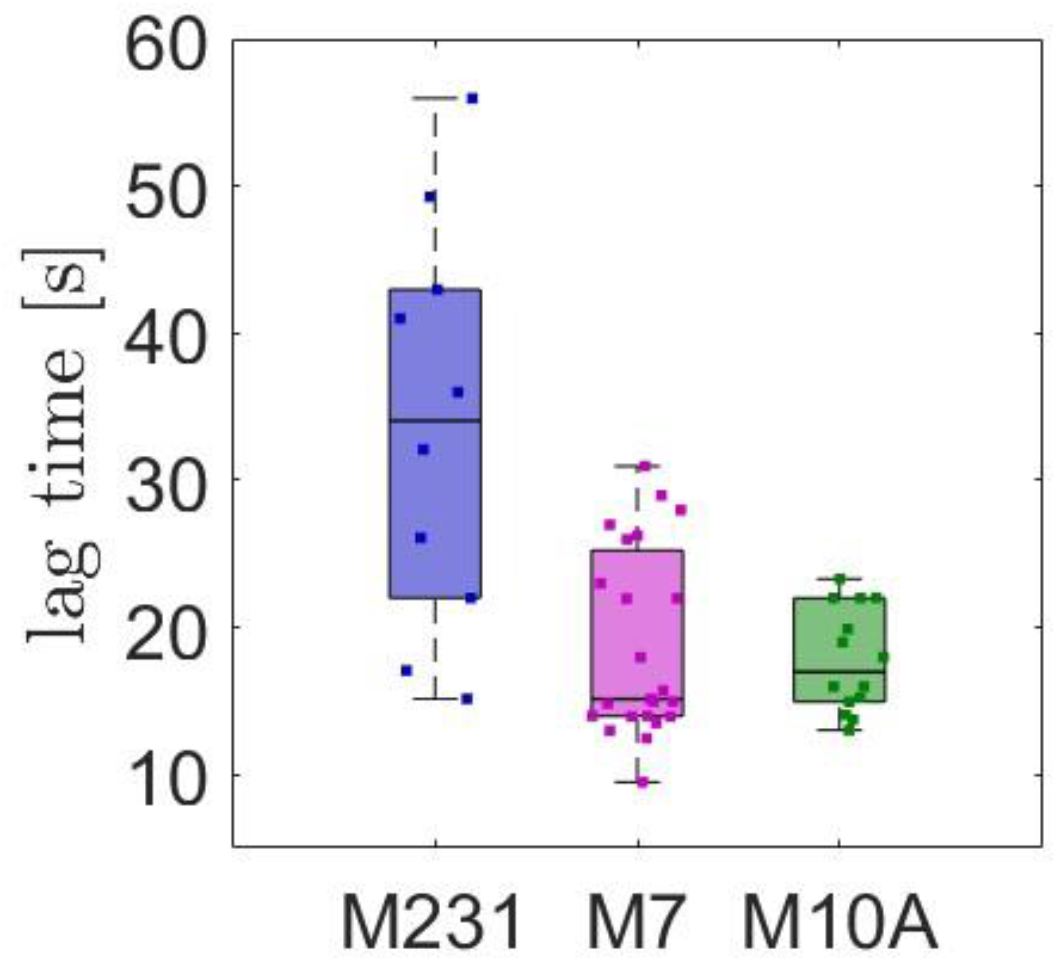
Lag time before relaxation monitoring. Box plots displaying the lag time for the three spheroids types. The lag time marks the time difference between the maximum strain experienced by the spheroid (before exiting the constriction channel) and the start of the relaxation recording. In other words, it is the time the spheroid takes to traverse the last serpentine and reach the relaxation chamber.

**Figure S2.**
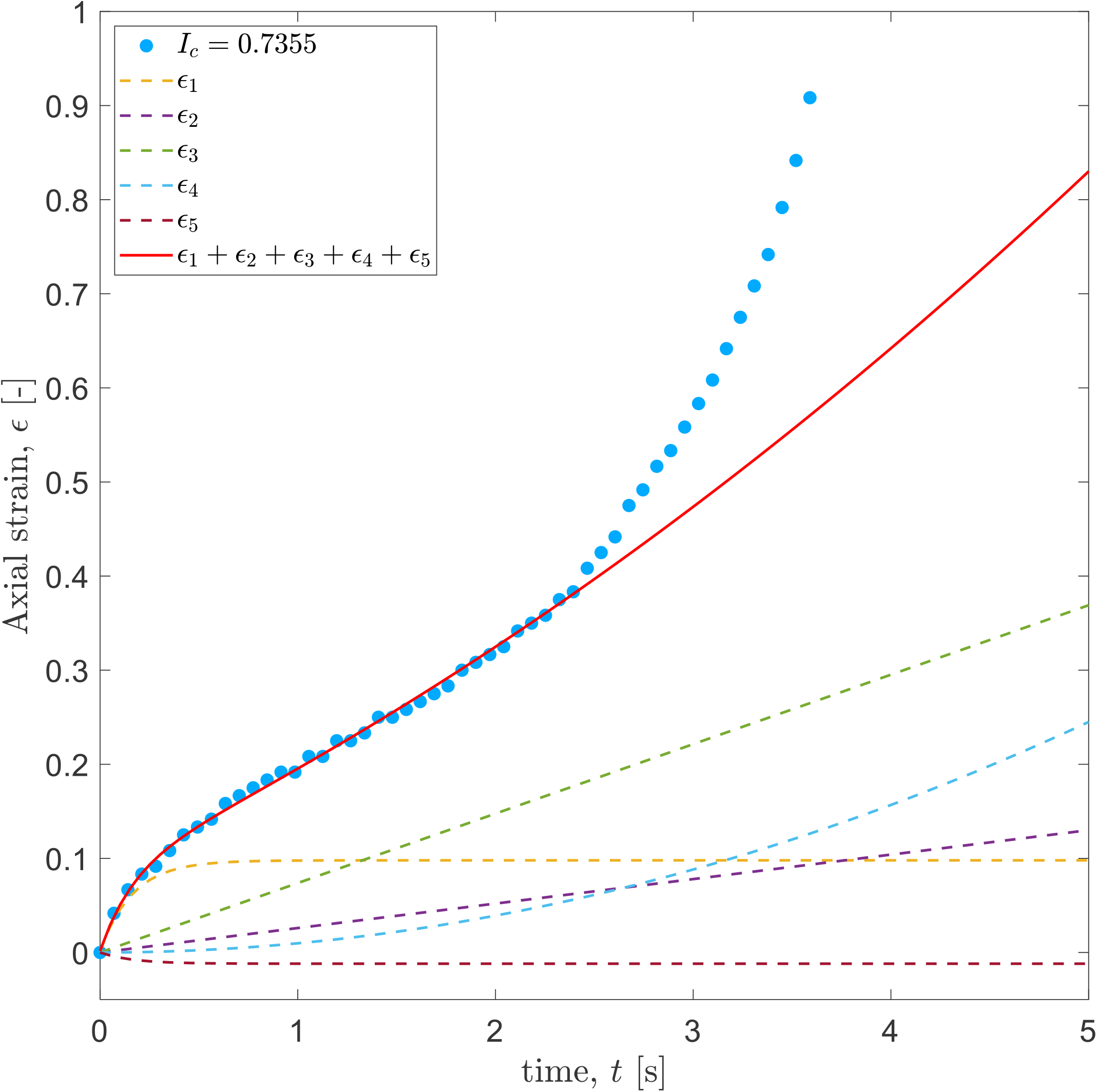
Decomposition of the constitutive equation of the D3M. 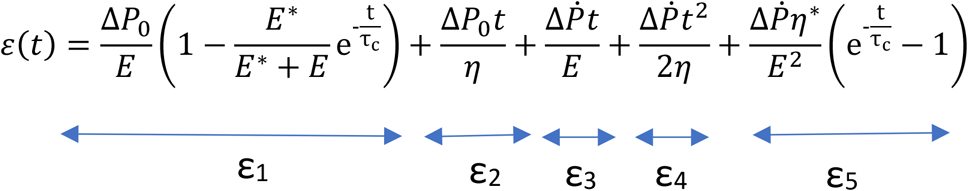

The red solid line shows the fitting of the D3M for a MCF-7 spheroid of *I*_*c*_ = 0.7355. The fitting stops at the inflex point, corresponding to the timepoint *t*_*2*_ as described in the main text. The dotted lines show the contribution of each term of the constitutive equation of the D3M, highlighting that the contribution of the last exponential term is very small

**Figure S3.**
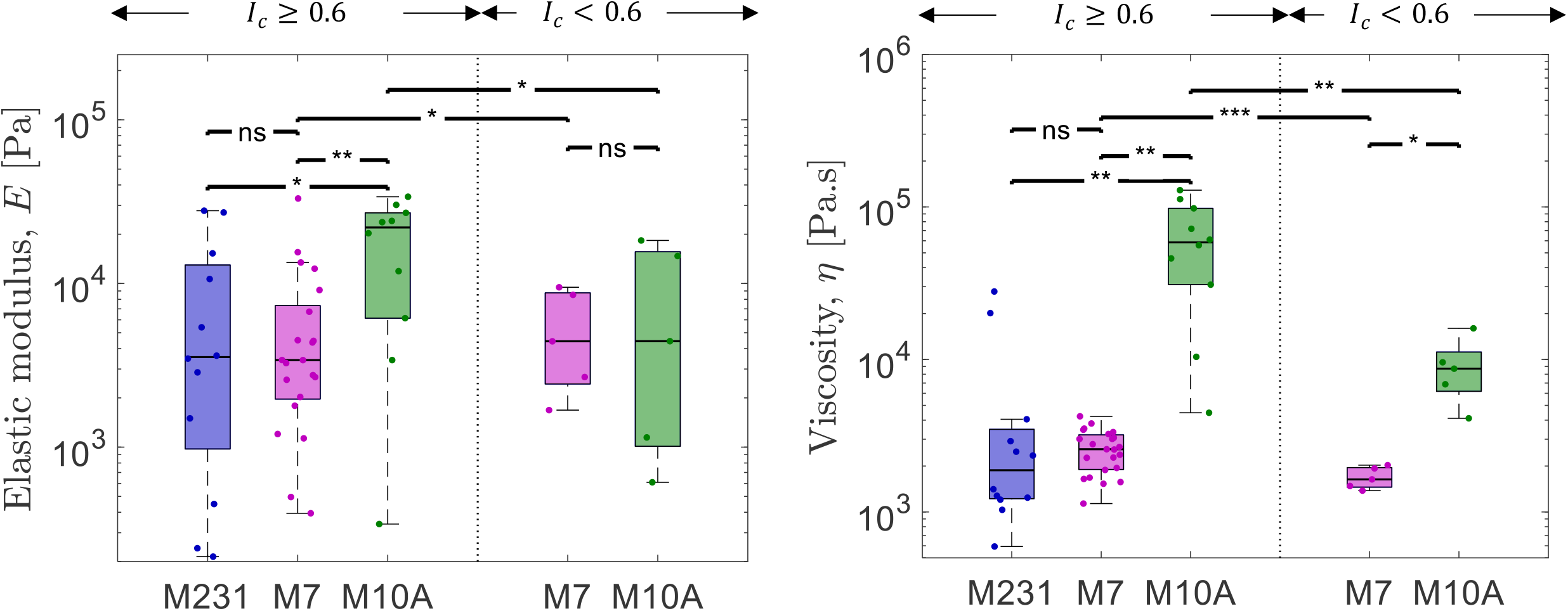
Spheroid viscoelastic properties extracted from 3M. Left: Bulk elastic moduli for the three spheroid types extracted from the fitting of the Modified Maxwell Model (3M). MCF-10A spheroids display higher bulk elasticity, compared to MCF-7 and MDA-MB-231 spheroids, with comparable values to the ones extracted from the D3M. Right: Viscosity for the three spheroid types extracted from the fitting of the Modified Maxwell Model (3M). The trend is similar to the one observed from the D3M, with MCF-10A displaying higher viscosity compared to the malignant spheroids.

**Table S1.**
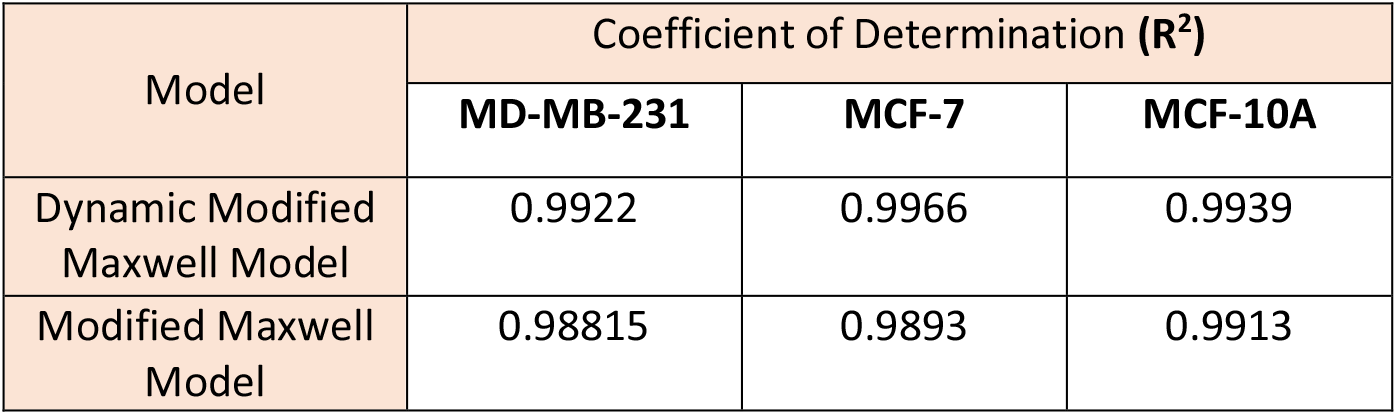
Comparison of the values of the coefficient of determination R^2^ for 3DM and 3M. The values reported represent the median value of the coefficient of determination R^2^ for all the fitting reported in both Figure 2e,f and Figure S4. The D3M shows a higher coefficient of determination compared to 3M for all three spheroid type.

**Figure S4.**
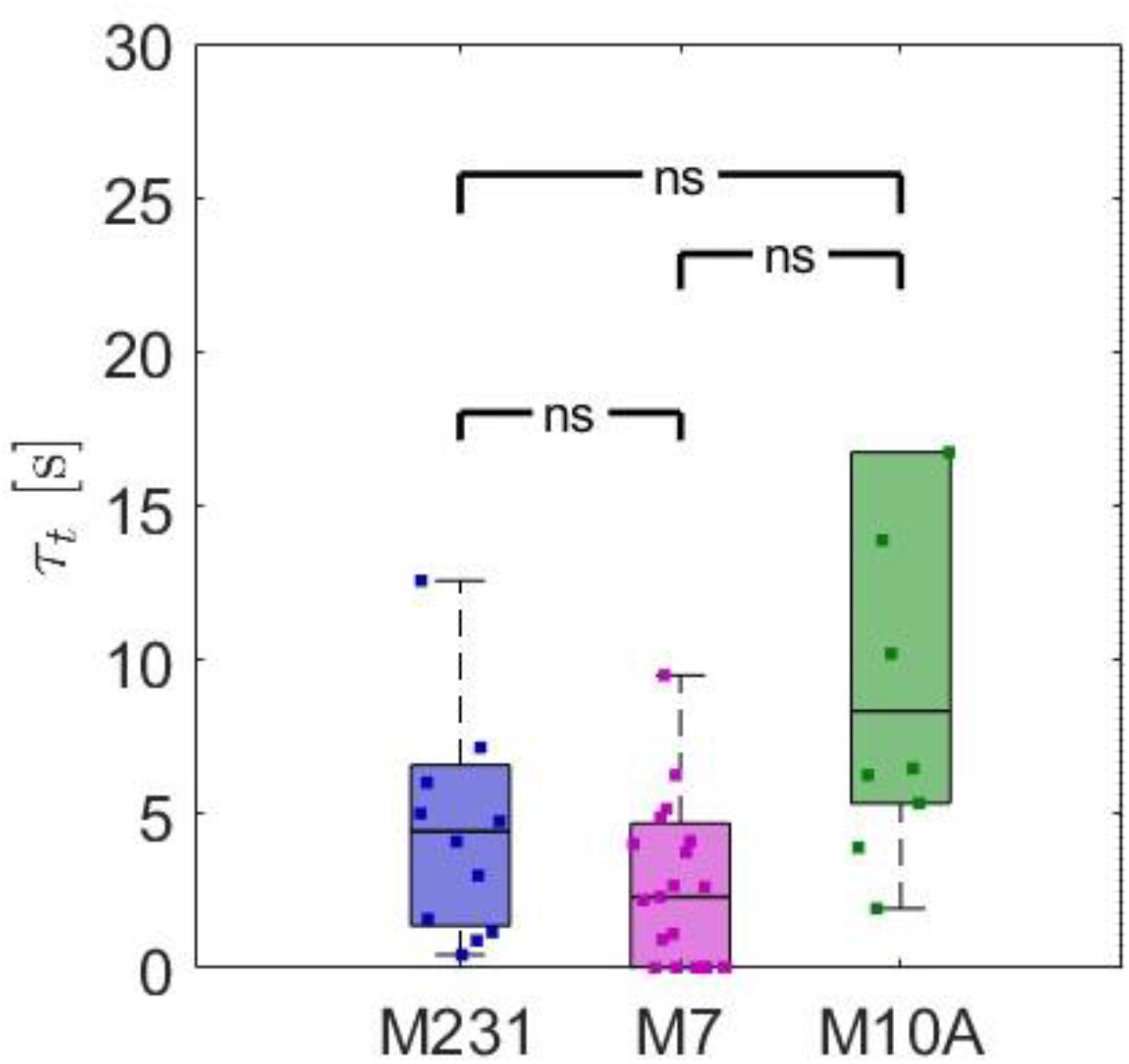
Tissue transition time. The transition time τ_t_, defined as η/E, describes the transition of the tissue from the elastic regime to the viscous regime during compression. MCF-10A spheroids show the longest transition time among the spheroids, suggesting a more predominant elastic contribution in the first stages of the dynamic compression in the benign spheroids.

**Figure S5.**
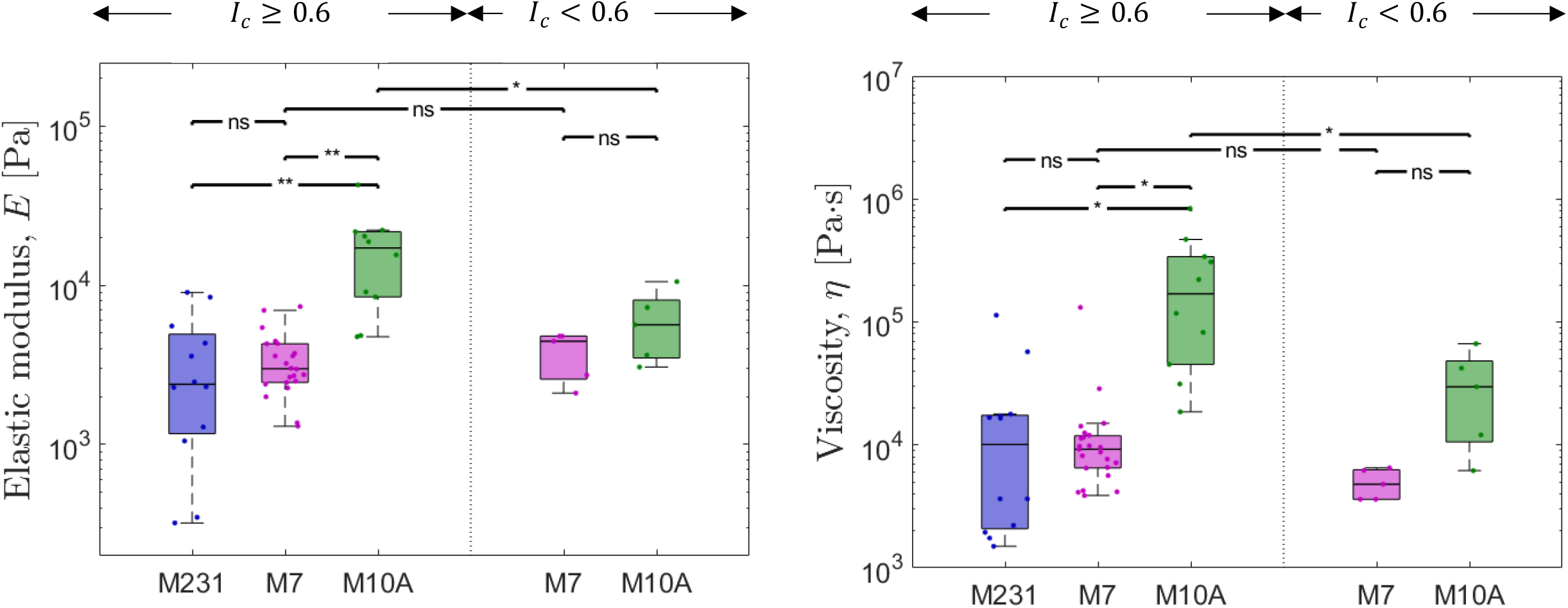
Bulk elastic moduli and viscosity for different I_c_ ranges. Bulk elastic moduli (left figure) and viscosity (right figure) for the three spheroid types for two ranges of I_c_. MCF-10A spheroids display viscoelastic properties that are dependent on the level of compression, with higher E and η as the compression increases (I_c_ ≥ 0.6). MCF-7 spheroids, instead, do not show any statistical differences in their viscoelastic properties across different I_c_ values. Interestingly, the distinction between benign and malignant spheroids disappears at lower levels of compression, indicating that the differences between them become discernible only above a certain threshold of stress or confinement.

**Figure S6.**
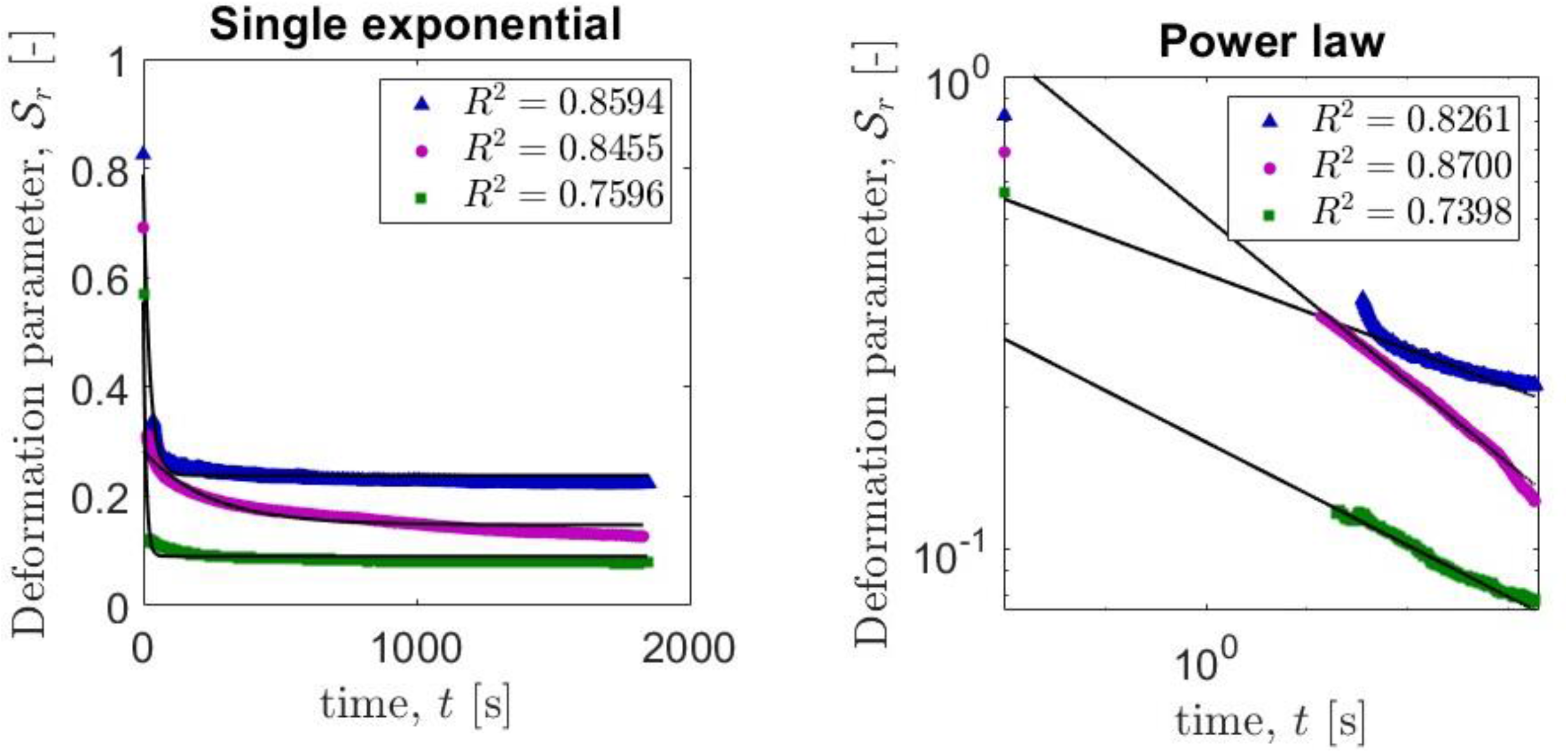
Relaxation curves fitting a single exponential and power law equations. Showcases of relaxation curves fitted with a single exponential (left) and power law (right, in log-log scale) equations. In the legends the coefficients of determination are outlined, highlighting that the double exponential model describes better the deformation parameter curves (with R^2^ > 0.98) via the two characteristic timescales.

**Figure S7.**
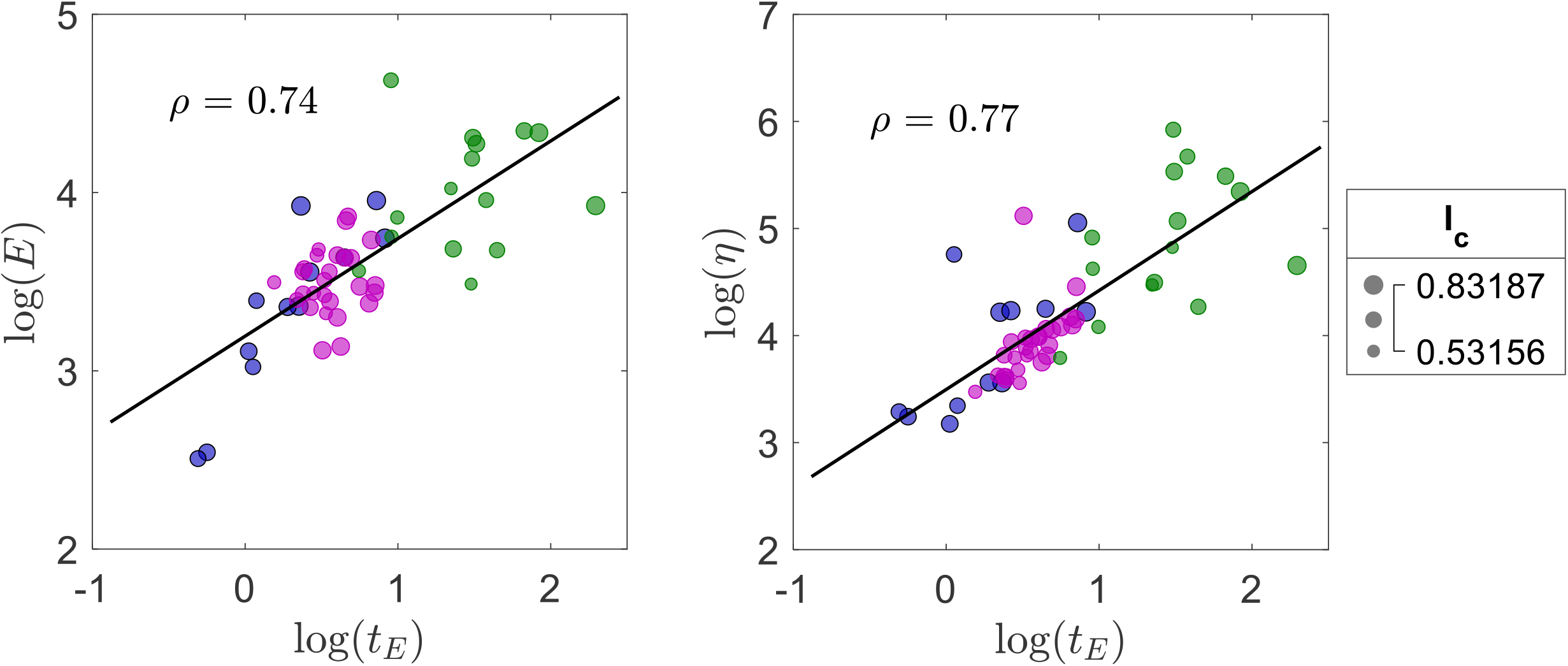
Bubble charts correlating entry time t_E_, bulk elasticity E viscosity η and Constriction Index I_c_. The colours correspond to the spheroid type: blue for MDA-MB-231,magenta for MCF-7 and green for MCF-10A. The bubble size increases with higher Constriction Index values. Left figure: Bubble chart showing the strong correlation (Pearson coefficient *ρ* = 0.74) between the entry time and the bulk elasticity of the aggregates, distinguished by spheroid type and I_c_ value. MCF-10A show higher bulk elasticity, corresponding to higher entry times, independently from the Constriction Index. Right figure: Bubble chart showing the strong correlation (*ρ* = 0.77) between the entry time and the aggregates viscosity, confirming the hypothesis that more viscous spheroids require longer time to deform and squeeze in the constricted channel. Smaller I_c_ values correspond to lower entry time t_E_ and for MCF-10A spheroids lower I_c_ translated in lower viscosity values, as shown in Figure S5.

## Supplementary Movies

**Movie S1**. Dynamic compression and deformation of MDA-MB-231, MCF-7 and MCF-10A spheroids respectively.

**Movie S2**. Relaxation of MDA-MB-231, MCF-7 and MCF-10A spheroids respectively.

**Movie S3**. Breakage and cell dissemination of MDA-MB-231 spheroid during dynamic compression and deformation

